# High-yield methods for accurate two-alternative visual psychophysics in head-fixed mice

**DOI:** 10.1101/051912

**Authors:** Christopher P Burgess, Armin Lak, Nicholas A Steinmetz, Peter Zatka-Haas, Charu Bai Reddy, Elina A K Jacobs, Jennifer F Linden, Joseph J Paton, Adam Ranson, Sofia Soares, Sylvia Schröder, Miles J Wells, Lauren E Wool, Kenneth D Harris, Matteo Carandini

## Abstract

Research in neuroscience relies increasingly on the mouse, a mammalian species that affords unparalleled genetic tractability and brain atlases. Here we introduce high-yield methods for probing mouse visual decisions. Mice are head-fixed, which facilitates repeatable visual stimulation, eye tracking, and brain access. They turn a steering wheel to make two-alternative choices, forced or unforced. Learning is rapid thanks to intuitive coupling of stimuli to wheel position. The mouse decisions deliver high-quality psychometric curves for detection and discrimination, and conform to the predictions of a simple probabilistic observer model. The task is readily paired with two-photon imaging of cortical activity. Optogenetic inactivation reveals that the task requires the visual cortex. Mice are motivated to perform the task by fluid reward or optogenetic stimulation of dopaminergic neurons. This stimulation elicits larger number of trials and faster learning. These methods provide a platform to accurately probe mouse vision and its neural basis.

## Introduction

The mouse is increasingly the species of choice for experiments that seek to understand the mammalian brain. Its advantages in ease of husbandry, breeding, and handling have been recognized for over 100 years, with the establishment of inbred lines that allowed researchers to control for genetic variation (Beck et al., 2000). Today the mouse offers an unrivaled arsenal of tools to the neuroscientist, from atlases of gene expression and connectivity (Lein et al., 2007; Oh et al., 2014; Zingg et al., 2014) to a vast array of genetic tools and transgenic lines (Harris et al., 2014; Heintz and Gerfen; Huang and Zeng, 2013; Madisen et al., 2015; Madisen et al., 2012). An additional advantage of the mouse is that its cortex is not folded, so it is more accessible to imaging studies.

Mice are also an excellent species for probing mechanisms of perception, decision, and cognition. Mice are readily trained to perform behavioral tasks based on touch (Guo et al., 2014a), olfaction (Liu et al., 2014; Resulaj and Rinberg, 2015), hearing (Hangya et al., 2015; Jaramillo and Zador, 2014; Pinto and Dan, 2015; Sanders and Kepecs, 2012) or vision (Andermann et al., 2010; Busse et al., 2011). Tasks based on vision, specifically, have been extended to probe not only sensation and perception, but also aspects of cognition (Bussey et al., 2012; Nithianantharajah et al., 2015).

Contrary to past preconceptions, mice make major use of vision (Carandini and Churchland, 2013; Huberman and Niell, 2011). The mouse visual cortex comprises a network of at least 12 retinotopic areas (Garrett et al., 2014; Glickfeld et al., 2014; Wang and Burkhalter, 2007). These areas might not map one-to-one to the 16-30 visual areas found in primates (Felleman and Van Essen, 1991; Markov et al., 2014; Wandell et al., 2007), but the principles governing the division of labor across areas are likely to be conserved across species (Wang et al., 2011). These and other principles of visual brain function may be fruitfully investigated in the mouse.

Studying the neural activity underlying visually-driven behavior, however, requires careful psychophysical techniques that pose specific constraints to the design of a task (Carandini and Churchland, 2013). (1) The task must allow continuous control of visual stimulation and accurate measurement of eye position. (2) It must be easily paired with brain recordings or manipulations. (3) It should require a behavioral response that does not confound the neural activity related to sensory stimulation and decision-making. (4) It should be robust to changes in the observer’s overall tendency to respond. (5) It should be learned relatively quickly and highly reliably by most subjects. (6) It should yield many trials per stimulus and per session, to deliver precise psychometric curves relating task performance to visibility. (7) It should yield close to 100% performance on easy trials, to distinguish errors due to the limits of vision from those that result from other sources (disengagement, confusion about the task rules, errors in motor control). (8) It should be flexible, so that its design can be made more complex if needed. (9) Finally, it would be ideal if one could motivate the subject to perform the task with purely positive rewards, without controlling their access to food or water.

These fundamental requirements are not met by the existing techniques for mouse visual psychophysics.

The first two requirements – careful control of visual stimulation and ability to perform brain recordings and manipulations – strongly argue in favor of head fixation, as some forms of brain recording or imaging can only be performed in an immobile brain. This requirement rules out techniques based on swimming (Prusky et al., 2000) or on poking the nose (Busse et al., 2011; Bussey et al., 2012; Long et al., 2015; Nithianantharajah et al., 2015). Some techniques available to study vision are compatible with head fixation, but they probe hard-wired subcortical behaviors such as the optokinetic reflex (Cahill and Nathans, 2008).

The third requirement – a behavioral response that does not confound sensory activity – rules out tasks where the behavioral report involves locomotion or navigation (Harvey et al., 2012; Poort et al., 2015; Wekselblatt et al., 2016). Locomotion and navigation elicit strong responses in mouse visual cortex (Niell and Stryker, 2010; Whitlock et al., 2008), and thus can confound sensory or decision-related signals that one may want to study there.

The fourth requirement – robustness to the observer’s tendency to respond – argues for having the observer choose between concurrent stimuli (Carandini and Churchland, 2013), as in a two-alternative choice design. This rules out go/No-go designs such as those where the mouse report the presence or absence of a visual stimulus by licking a single spout (Andermann et al., 2010; Glickfeld et al., 2013; Goard et al., 2016; Lee et al., 2012). Promising methods for two-alternative choices in head-fixed mice are available to probe audition, somatosensation, and olfaction (Guo et al., 2014a; Resulaj and Rinberg, 2015; Sanders and Kepecs, 2012) but have not been adapted to probe vision. Finally, no existing techniques meet the ideal requirement of a positive reward with no implicit punishment. In current tasks, the reward simply redresses an unpleasant circumstance: swimming in deep water (Prusky et al., 2000), or having limited access to drinking water (Andermann et al., 2010; Busse et al., 2011; Bussey et al., 2012; Glickfeld et al., 2013; Lee et al., 2012; Long et al., 2015; Nithianantharajah et al., 2015).

We thus developed a task that meets all the above requirements, where the behavioral response is based on a steering wheel. In choosing this manipulandum we were inspired by tasks developed to probe hearing and olfaction, where mice use their front paws to rotate a conveyor belt or a spherical ball (Resulaj and Rinberg, 2015; Sanders and Kepecs, 2012). In our task, mice use their front paws to turn the steering wheel left or right, providing a two-alternative choice between concurrent visual stimuli. To train the mice in this task, we found it useful to introduce an intuitive coupling of the steering wheel to the position of the visual stimuli. With this coupling, mice learn this task within a few weeks, they perform it proficiently, and their decisions conform to the predictions of a simple probabilistic observer model. The task can be readily paired with two-photon imaging, activates visual cortex, requires visual cortex, and can be flexibly extended to probe unforced-choice, both for stimulus detection and for stimulus discrimination.

Mice performed the task not only when rewarded with water but also when rewarded with selective stimulation of midbrain dopaminergic neurons. Optogenetic stimulation of these neurons is known to elicit coarse behavioral outcomes such as place preference (Tsai et al., 2009) or simple repetitive actions such as nose-poking (Kim et al., 2012). Here we show that it acts as a powerful reward in precise actions driven by perceptual decisions.

## Results

We begin by introducing the basic design of the task: two-alternative forced-choice contrast detection with a water reward, and we show that this task is compatible with precise recordings of visual responses in cortex. We then introduce the unforced-choice version of the task, we define a simple probabilistic observer model for the mouse decisions, and we show how these decisions are impaired by optogenetic inactivation of visual cortex. Finally, we illustrate two variations of the method: one where the reward is optogenetic stimulation of dopaminergic neurons rather than water, and one where the task involves discrimination between two stimuli in opposite visual fields.

### The basic task: two alternative forced-choice

To allow a head-fixed mouse to select one of two choices, we placed a steering wheel under the front paws, and coupled its angular position to the position of a visual stimulus on the screen (Figure 1**a**). We chose the wheel as manipulandum (Figure 1**a**, *left*) as it resembles those successfully used to probe mouse audition (Sanders and Kepecs, 2012) and mouse olfaction (Resulaj and Rinberg, 2015). To train mice to use this manipulandum in a visual task, it was highly advantageous to couple wheel movements to the visual stimuli, so that turning the wheel left or right would accordingly move the stimuli left or right (Figure 1**a**, *right*; Supplementary Movie 1). The mouse indicates its choice of stimulus by bringing the stimulus to the center of the visual field.

The typical sequence of trial events was as follows (Figure 1**b**). First, the mouse had to keep the wheel still to initiate the trial and make the stimulus appear. Second, an Onset tone signaled the arrival of the stimuli, and an “open loop” period began, during which wheel movements were ignored. Mice generally continued to hold the wheel still in this period, and if the experiment required this behavior, it was reinforced through training. Third, a Go tone was typically played (typically, a 12 kHz pure tone lasting 100 ms with a 10 ms onset and offset ramp), after which the mouse could respond at any time, and wheel turns resulted in movements of the visual stimuli (“closed loop”, Figure 1**b**). If the mouse turned the wheel such that the stimulus reached the center of the screen, the stimulus locked in place and the animal received a small amount of water (1-3 µL). If instead the mouse moved the stimulus by the same distance but in the opposite direction, the stimulus would lock in place there and this incorrect decision was penalized with a timeout (typically, 2 s) signaled by auditory noise. In either case, the grating typically remained locked in its response position for 1 s to remind the mouse of its action while it received its feedback, and then disappeared.

Depending on the requirements of the specific experiments, in many mice we adopted slight variations of this sequence of events. For instance, if an experiment could tolerate motor actions prior to visual stimulation, we dropped the requirement for a quiescent period. Similarly, we introduced the open loop period only if we wanted to avoid motor actions or visual motion during stimulus presentation. Likewise, we played the Onset and Go tones only if we did not mind evoking auditory activity, and we shortened the inter-trial interval when we were striving for a larger number of trials in a session. Multiple variations of this task are thus possible. Our analyses here do not distinguish among these variations because other key factors covaried with them: experimenter, time of day, experimental rig, home cage, etc. A proper comparison would have to correct for these factors, and we do not have the data to do so.

**Figure 1.**
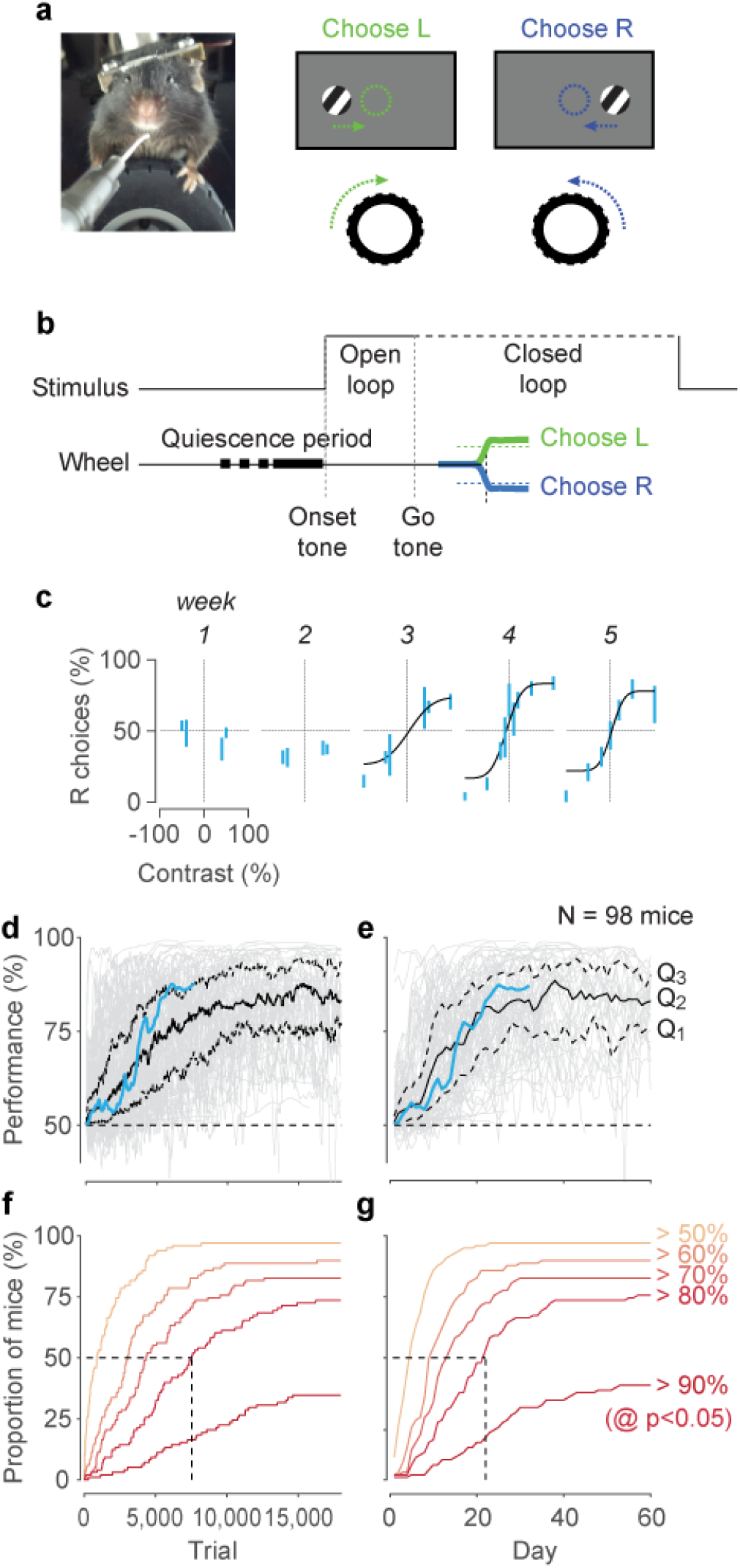
Fundamentals of the stimulus detection task, in its basic two-alternative forced-choice (2AFC) version. **a**. Left: A typical setup showing the head-fixed mouse with forepaws on a steering wheel used to make choices. Right: Schematic of the two possible stimulus conditions. At the onset, the grating is either on the left or on the right, and the mouse must turn the wheel (arrows) to move the grating to the final rewarded position (dashed circles). **b**. Time course of the basic task. Mice start the trial by holding the wheel still (quiescence). An onset tone may be played. The stimulus appears. Its position is initially fixed, i.e. it cannot be moved by moving the wheel (open loop). After an optional Go tone, stimuli become paired with wheel position (closed loop). Choices are made when the stimulus reaches the center of the screen (this choice is rewarded) or an equal distance in the opposite direction (this choice is penalized with a timeout and a noise sound). **c**. Psychometric data obtained in the first five weeks for an example mouse. Bars show the percentage of times the mouse chose the right stimulus (95% binomial confidence intervals), as a function of stimulus contrast. By convention we plot contrast of stimuli on the left as negative, and contrast on the right as positive. In later weeks, the data are fitted with a psychometric curve. **d**. Learning rates for a population of 98 mice. Performance is assessed on highly visible stimuli (≥40% contrast), as a function of number of trials. Blue trace highlights the example mouse in **c**. Gray traces indicate performance by individual mice, with black traces indicating the three quartiles: the median (Q2) and the 25th and 75th percentiles (Q1 and Q3). The approximate chance level is 50% (dashed line). These 98 mice were trained in the basic version of the task and in slight variations of the task. **e**. Same as **d** but expressed as a function of training days rather than trials. **f**. Cumulative probability curves showing the proportion of mice surpassing a given performance level as a function of trial number, with a significance level of p<0.05. **g**. Same as **f**, but expressed as a function of training days rather than trials.

Training for a typical mouse proceeded in two main stages (Figure 1**c**-**e**). We started the mouse on easy (high) contrasts, until it learned the association between turning the wheel and moving the stimulus, and receiving rewards. In our experience, this association (the closed loop period) is necessary for learning: in a few attempts where we did not use the closed loop period, the mice did not learn the task. When the mouse was above chance level performance for a day or two (which typically occurred by the first week), we began to introduce lower contrasts. For example, a typical mouse (Figure 1**c**) reached 56% performance (within a 95% binomial confidence interval) on a high contrast stimulus after ~2,300 trials (Figure 1**d**, *blue*), on day 5 (Figure 1**e**, *blue*), after which we introduced lower contrast stimuli. Psychometric functions of stimulus contrast and position were obtained by week three (Figure 1**c**). By week four, this animal had mastered the task.

These results were typical of our population (n = 98 mice, Figure 1**d**-**g**). Most mice were above chance before ~1,000 trials (Figure 1**d**), corresponding to a few days of training (Figure 1**e**). Mice then typically approached steady performance after 7,000-10,000 trials (Figure 1**d**) i.e. in 20-30 days (**e**). Only few mice (6/98) failed to learn the rudiments of the task (performance significantly above 50%) by trial 5,000 or after two weeks (Figure 1**f,g**). Most animals surpassed 80% performance, but a sizeable fraction (38/98) also reached 90% performance (Figure 1**f,g**). The method therefore worked in practically all mice, even though different cohorts were trained by different experimenters using different subjective criteria about when to advance a mouse from one stage of training to the next.

Once they mastered the task, mice typically produced consistent movements, with initial wheel deflections usually matching the final responses made (Supplementary Figure 1). The movements showed some dependence on stimulus attribute (responses to stimuli with higher contrast typically had shorter latency and higher peak velocity) but were otherwise fairly stereotyped, showing little variability across trials. If desired, we could then modify the task by removing the coupling between wheel position and stimulus position, so that the stimulus would stay fixed in its position (Supplementary Figure 2), or disappear as soon as the movement started.

Some mice tended to move their eyes following stimulus onset, or showed changes in pupil diameter associated with the trial structure (Supplementary Figure 3). These eye movements and pupil dilations, however, were highly variable across trials and across mice, highlighting the importance of imaging the eye in all experiments.

### Simultaneous recordings in visual cortex

To confirm that this task could be readily paired with measurements of brain activity, we performed two-photon imaging of activity in primary visual cortex (V1) of mice that were performing the task (Figure 2). We expressed GCaMP6m in V1 neurons (right hemisphere) via AAV2/1 virus injection, and trained the mice in a version of the task where trials started only after the mouse had held the wheel still for a 2-3 s quiescence period, there was no Go tone, and the “open loop” period lasted 1 s (Figure 1**b**). During this period we could image neural responses without the stimulus moving. While mice performed this task (Figure 2**a**) we performed two-photon calcium imaging of V1 neurons, choosing a field of view with cells whose receptive field overlapped with the (contralateral) stimulus (Figure 2**b**).

As expected, most visually-responsive cells showed robust responses to contralateral stimuli, and gave no response to ipsilateral stimuli (Figure 2**c**-**d**). The onset of contralateral stimuli evoked strong calcium transients during the open loop period (Figure 2**c**). The amplitudes of these transients grew with the contrast *c* of contralateral gratings (Figure 2**d**). We fit these responses with the commonly used function

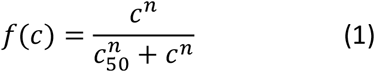

where *c*_50_ and *n* are free parameters (Albrecht and Hamilton, 1982; Sclar et al., 1990). Similar results were routinely obtained in other mice (e.g. Figure 2**e**-**h**). These results demonstrate that the visual task can be readily paired with recording techniques that require high stability, and that it evokes contrast-dependent activity in cortex.

Moreover, these data indicate that movements of the steering wheel did not by themselves cause overwhelming activation in visual cortex. Indeed, there was little if any activity when high-contrast stimuli (which almost invariably caused movement) were presented ipsilaterally.

This said, V1 activity did include small fluctuations that tended to precede wheel movements (Figure 2i**-k**). The traces from three of the example neurons illustrate this phenomenon: activity was not confined to the large responses elicited by contralateral stimuli (positive contrasts, Figure 2i). There were additional fluctuations, whose timing related to the movements of the wheel (Burgess, 2016). Triggering these small fluctuations on the onset of the wheel turns (in the absence of visual stimuli) shows that activity typically built up during periods of wheel quiescence, perhaps reflecting increased alertness, and decayed following the onset of wheel movements (Figure 2j**-k**). This buildup of activity, however, was generally dwarfed by visual responses. For instance, for the 6 example cells (Figure 2**b-d,f-h**), the activity measured during the build-up was 7.5 ± 0.8 times smaller (mean ± s.e.) than the responses to 50% contrast ipsilateral visual stimuli.

**Figure 2.**
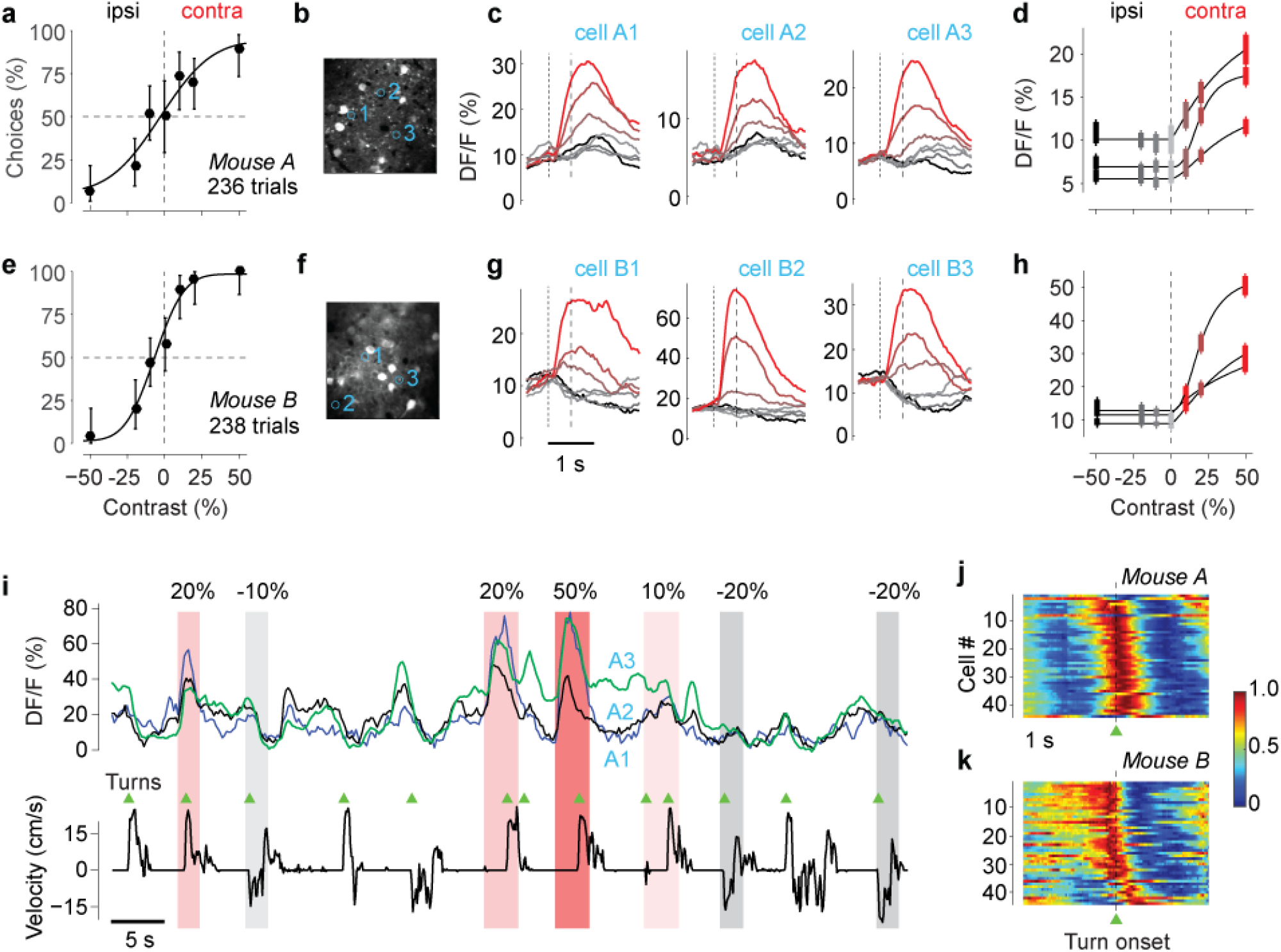
Imaging in V1 during the task. **a**. Psychometric curve for an example mouse, measured during two-photon imaging in area V1. **b.** Imaging field of view, with three cells (regions of interest) circled and numbered. **c**. Mean calcium activity averaged around the onset of the grating stimulus, grouped by stimulus condition (color codes are given in the next panel) for the three cells. Dotted line indicates stimulus onset (preceded by 2-3 s quiescence period). Dashed line indicates the beginning of the interactive period, when the stimulus becomes movable (end of open loop). These data were taken from 181 trials (22-30 per condition). **d**. Response amplitudes of each cell as a function of stimulus contrast. Positive contrast denotes stimuli in the contralateral visual field, and negative contrast denotes stimuli in the ipsilateral visual field. Amplitude is mean response at 1 s after grating onset. Curves indicates fits of the function *p* + *qf*(*c*), with *f*(*c*) defined in Equation 1. Error bars indicate s.e.m. **e**-**h**. Same as **a**-**h**, for a different mouse. Data were taken from 210 trials (24-43 per condition). **i**. Example traces from the three cells in **b**-**d**, in the presence of stimuli of different contrasts (shaded areas) and in relation to wheel velocity (*bottom trace*). There are strong responses to the visual stimuli but also small responses synchronized with turn onsets (*triangles*). Onsets and offsets of wheel turns were identified by applying a dynamic threshold procedure based on a Schmitt trigger to the wheel velocity traces. **j-k.** Time course of movement-related activity (in the absence of visual stimuli) in 45 neurons from each of the two mice. These cells were the top ranking cells in each experiment, based on the quality of segmentation. We triggered calcium activity on wheel turn onsets, averaged across events, and normalized the results for each neuron (rows) to range from 0 to 1. Neurons are sorted by the amplitude 1 s before turn offset.

### Two alternative unforced-choice (2AUC)

In many conditions, it helps to extend a two-alternative task by allowing a “no-go” response option when the stimulus is absent. With this two-alternative unforced choice (2AUC) version of the task, one can measure sensitivity and bias separately for the two stimulus locations. This is particularly useful following unilateral manipulations in task context or brain activity (Sridharan et al., 2014).

We trained mice in a 2AUC version of the contrast detection task and found that they were readily able to perform it (Figure 3**a**-**c**). After training mice on the 2AFC task, we added the no-go condition: when the stimulus was absent (zero contrast), mice would earn the reward by refraining from turning the wheel (no-go, Figure 3**a**) for the duration of a 1.5 second response window (Figure 3**b**). Mice typically learned this new response contingency in 5-6 sessions. Their reaction times for responses left or right were much faster than the 1.5 s response window (Figure 3**b**, and Supplementary Figure 4), indicating that issuing a No-go response was distinct from simply being slow to respond. Consistent with this interpretation, mice made most No-go choices at zero contrast (when these choices were correct), and made progressively fewer of them as stimulus contrast increased (Figure 3**c**).

**Figure 3.**
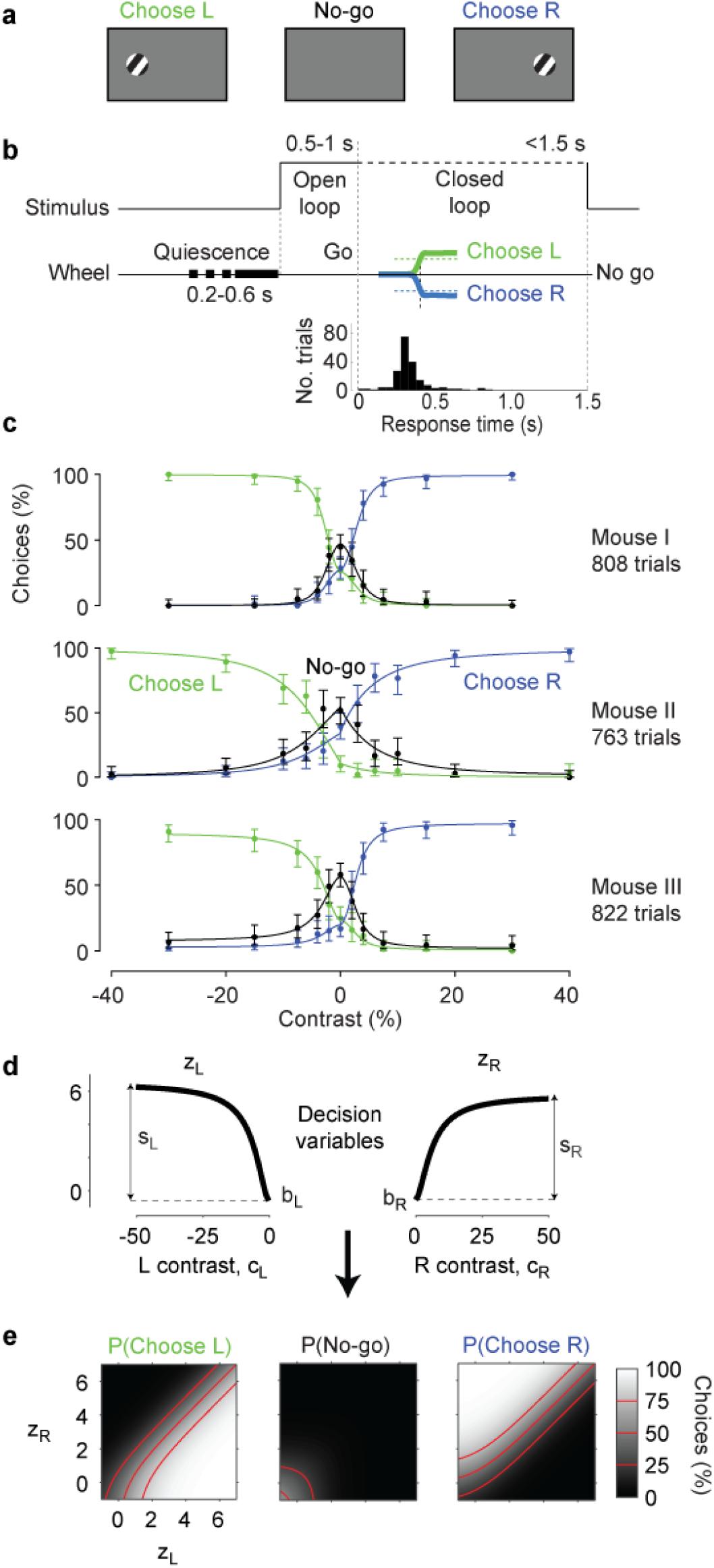
Elaboration of the stimulus detection task in a two-alternative unforced-choice (2AUC) version. **a**. In the 2AUC task, the mouse learns to choose left when the stimulus is on the left, choose right when the stimulus is on the right, and make neither response (No-go) if the stimulus is absent. **b**. Time course of the 2AUC task. At the Go cue, the mouse has 1.5 s to move the wheel. Holding the wheel still for this period counts as a No-go choice, which his rewarded if the stimulus was absent (zero contrast). *Histogram* shows a typical distribution of response times in a session (time from go tone to completion of wheel turn). **c.** Choices as a function of stimulus contrast and position for three individual sessions in three mice (rows). For each mouse the data show the proportion of left choices (green), right choices (blue), and no-go choices (black), as a function of stimulus contrast. As in all other plots, negative contrast denotes stimuli appearing on the left side. The data were fitted with the probabilistic observer model (smooth curves). Error bars are 95% binomial confidence intervals. **d**. The decision variables in the probabilistic observer model, with parameters obtained from the first mouse in **c**. The decision variables z_L_ and z_R_ grow as a function of contrast presented on the left and on the right, respectively. These functions are each defined by two parameters, a bias *b* and a sensitivity *s* (Equations 1 and 2). **e**. The probability of each of the three possible choices (Left, No-go, and Right) depends on the two decision variables. This dependence is fixed, i.e. parameter-free (Equation 3).

This 2AUC version of the task thus yields three psychometric curves: one for the probability of choosing L, one for the probability of choosing R, and one for the probability of No-go (Figure 3**c**). This representation is redundant, because the three probabilities must sum to 1. In other words, only two of the three curves are independent, and the third is fully constrained from the other two. Nonetheless, it helps to view all three to understand the data, and it helps to develop a simple observer model to interpret the data and fit them. We introduce this model next.

### Probabilistic observer model

The decisions made by the mice in the task conformed closely to the predictions of a simple probabilistic observer model. We present here the model for the 2AUC version of the task, because it is the more general case; the model can be easily reduced to the 2AFC version of the task.

In the model, choices depend on two decision variables, one for choosing L and one for choosing R (Figure 3**d**). These two decision variables, *z*_*L*_ and *z*_*R*_, grow with the contrast on the left *c*_*L*_ and on the right *c*_*R*_ according to a simple expression:

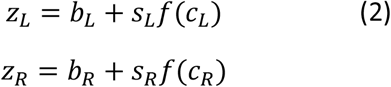

Here, the parameters *b*_*L*_ and *b*_*R*_ measure the bias towards choosing left or right (relative to choosing No-go), the parameters *s*_*L*_ and *s*_*R*_ measure the weight assigned to visual evidence on the left or right, and *f*(*c*) is the function of contrast in Equation 1 (Figure 3**d**).

The decision variables, in turn, determine the probability of choosing L or R, and thereby, the probability of choosing no-go (Figure 3**e**). Specifically, the decision variables are identical to the log odds of choosing left or right vs. choosing No-go:

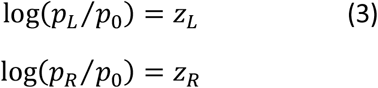

where *p*_*L*_ and *p*_*R*_ are probabilities of choosing L or R, and *p*_0_ is the probability of choosing No-go. These two expressions are parameter-free. They fully describe the three probabilities because *p*_*R*_ + *p*_*L*_ + *p*_0_ = 1.

**Figure 4.**
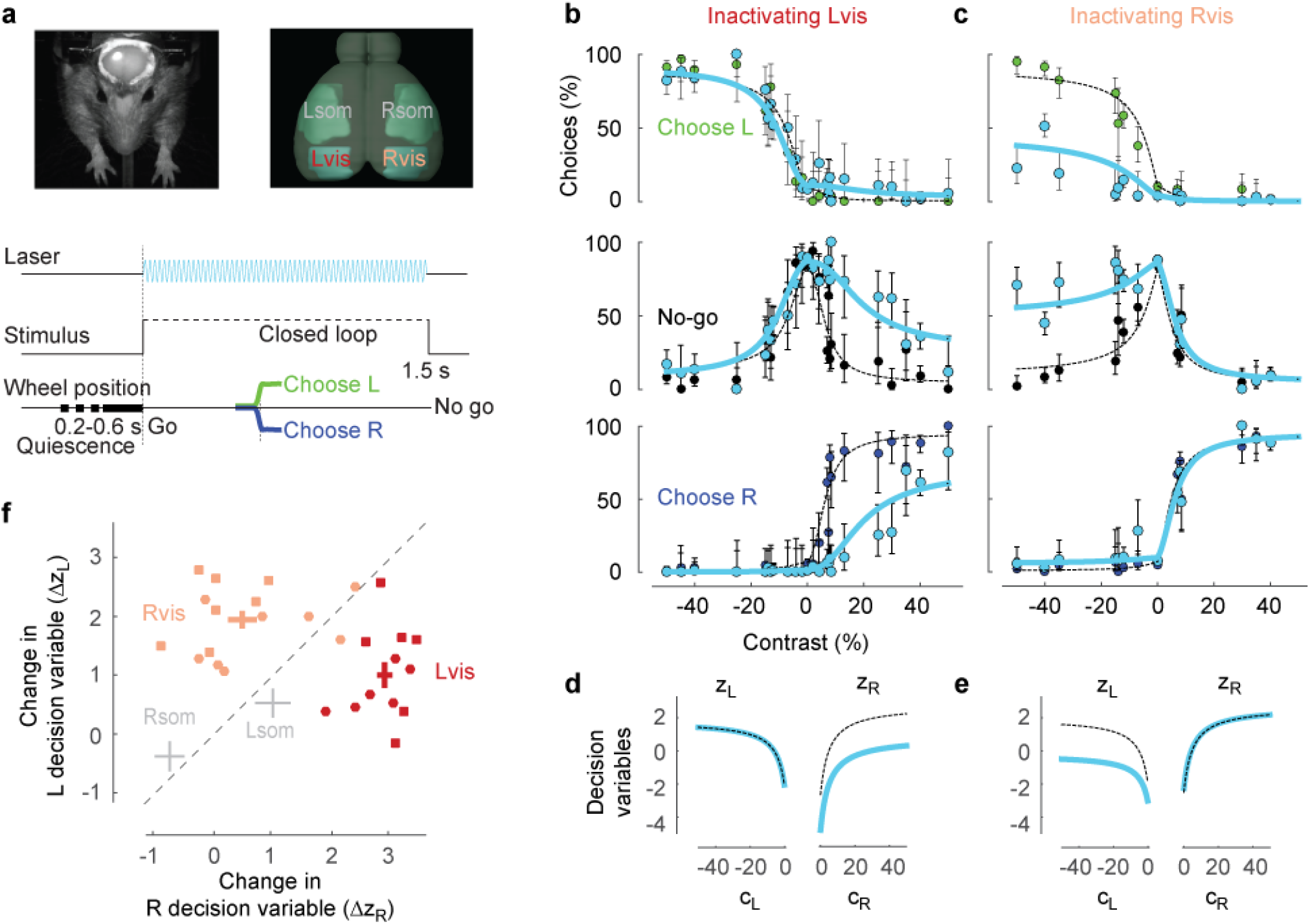
Effects of optogenetic inactivation of visual cortex. **a**. Methods of optogenetic inactivation during the 2AUC task. *Top left*: image of a mouse with the clear skull preparation, with laser spot on right hemisphere. *Top right*: illustration of the two regions inactivated in each hemisphere: left and right visual cortex (Lvis and Rvis) and, as a control, left and right somatosensory cortex (Lsom and Rsom). Inactivation of these regions was performed in different sessions. *Bottom*: Time course of the task (this variant did not include an open loop period). On ~33% of trials, stimuli were accompanied by laser illumination. **b**. Effects of inactivation of left visual cortex. Proportion of left, no-go, and right choices as a function of stimulus contrast, under control conditions (green, black, and blue dots) and during optogenetic inactivation (cyan dots). Curves indicate fits of the probabilistic model under control conditions (dashed) and during optogenetic inactivation (cyan). Error bars show 95% binomial confidence intervals. Data were obtained in 6 sessions from one mouse (2,521 trials). **c**. Same as **b**, for inactivation of right visual cortex from the same mouse. Data were obtained in 7 sessions (2,782 trials). **d**. Decision variables obtained by the model fits in **b**, as a function of contrast on the left and right, in control condition (dashed) or during inactivation of left visual cortex (cyan). **e**. Same as **d**, for inactivation of right visual cortex. **f**. Summary of the effects of optogenetic inactivation in the four regions outlined in **a**. Effects are measured by the decrease in the left and right decision variables *z*_*L*_ or *z*_*R*_ at 50% contrast. Dots indicate individual sessions from two mice (squares for the mouse in **b**-**e**, circles for another mouse) with inactivation of left visual cortex (red) or right visual cortex (pink). Crosses summarize the effects of inactivation in visual cortex (red and pink), and in somatosensory cortex (gray). The length of the crosses indicates ± s.e.m. in the two dimensions.

With 6 free parameters, the model provided good fits to the 22 measurements, explaining over 75% of individual choices (curves in Figure 3**c**). The only free parameters in the model are those that relate the two contrasts to the two decision variables (Figure 3**d**): 2 parameters in Equation 1, and 4 parameters in Equation 2. Fitting procedures and model parameters are described in Methods. Cross-validation indicated that for these three data sets there would be no loss in fit quality if one imposed *s*_*L*_ = *s*_*R*_, thus removing one free parameter. However, as we will see next, these two parameters must be allowed to be different when evaluating the effects of manipulations such as unilateral inactivation.

### Inactivation in visual cortex

To assess the involvement of visual cortex in task performance, we employed optogenetic techniques to inactivate cortical areas during individual trials (Figure 4**a**). We used two transgenic mice expressing ChR2 in *Pvalb*-positive inhibitory interneurons and implanted them with clear skull caps (Guo et al., 2014b). We use a 473 nm laser to inactivate a cortical region during visual stimulus presentation and wheel-turn responses. Electrophysiological measurements show that such inactivation was circumscribed to a radius of ~1 mm (Supplementary Figure 5). We centered the inactivation on the left or right visual cortex or, for control measurements, on the left or right somatosensory cortex.

Inactivation of visual cortex strongly suppressed the mouse’s ability to detect contralateral stimuli (Figure 4**b,c**). Inactivating left visual cortex had barely any effect on the detection of ipsilateral stimuli, but greatly increased the fraction of no-go responses to contralateral stimuli, and correspondingly decreased the correct detection of those stimuli (Figure 4**b**). Analogous results were seen when inactivating visual cortex in the right hemisphere (Figure 4**c**).

To summarize these effects and compare them across experiments, we used the probabilistic observer model (Figure 4**d**-**f**). In the example experiment, inactivation of left visual cortex reduced only the decision variable for right stimuli (z_R_, Figure 4**d**), and inactivation of right visual cortex reduced only the decision variable for left stimuli (z_L_, Figure 4**e**). Similar results were seen across experiments (Figure 4**f**): inactivation of left visual cortex decreased z_R_ by 2.9±0.1, significantly more than z_L_ (1.0±0.2; paired t-test, one-sided, p<10^-5^), and inactivation of right visual cortex decreased z_L_ by 2.0±0.2, significantly more than z_R_ (0.5±0.2; p<10^-4^).

By comparison, in control experiments where we inactivated the somatosensory cortex we saw no such effects (Figure 4**f**, Supplementary Figure 6). Inactivation of somatosensory cortex did not cause any significant change in decision variables (p=0.17 and p=0.25 for inactivation of left and right somatosensory cortex, Supplementary Figure 6). Indeed, the effect of inactivation on decision variables was significantly weaker in somatosensory than in visual cortex. The effect on the R decision variable was significantly weaker during inactivation of left somatosensory cortex than of left visual cortex (p=0.00015, Wilcoxon rank sum test). Similar effects were seen on the L decision variable following inactivation of right somatosensory vs. visual cortex (p=0.00012). We conclude that accurate performance on this task requires that mice make specific use of their visual cortex.

### Optogenetic dopamine reward

The conventional method to reward mice for performing perceptual decisions involves delivering fluids under conditions of water control (Andermann et al., 2010; Busse et al., 2011; Bussey et al., 2012; Carandini and Churchland, 2013; Glickfeld et al., 2013; Guo et al., 2014a; Lee et al., 2012; Long et al., 2015; Nithianantharajah et al., 2015). The purpose of water control, however, is simply to make water rewarding. It would be ideal if one could deliver rewards without any water or food control. We sought to achieve this goal by providing direct stimulation of the brain centers that mediate the effects of positive reinforcement.

To motivate the animals to learn and perform the task, we provided phasic optogenetic stimulation of midbrain dopamine (DA) neurons. Phasic stimulation of these neurons, delivered electrically or optogenetically, is known to be sufficient for simple behavioral conditioning, such as place preference or lever pressing or nose poking (Kim et al., 2012; Olds and Milner, 1954; Tsai et al., 2009). However, it is not known whether trial-by-trial stimulation of these neurons could act as an efficient reinforcement for training a perceptual choice task.

We injected a viral construct containing Cre-dependent Channelrhodopsin-2 (ChR2) into ventral tegmental area (VTA) and substantia nigra pars compacta (SNc) of DAT^IREScre^ mice, and we implanted an optic fiber above VTA (Figure 5**a**). We confirmed specific expression of ChR2 in dopamine neurons using immunohistochemistry (Figure 5**b**). We identified dopaminergic neurons as those that stained for tyrosine hydroxylase (TH+). 71% of these neurons also expressed ChR2. On the other hand, only 5% of neurons that expressed ChR2 failed to react to TH staining, indicating that the expression was highly selective to DA neurons. The expression of ChR2 was consistent across animals and was highly stable up to 7 months (our last measured data point) after virus injection (n = 11 mice, Supplementary Figure 7).

**Figure 5.**
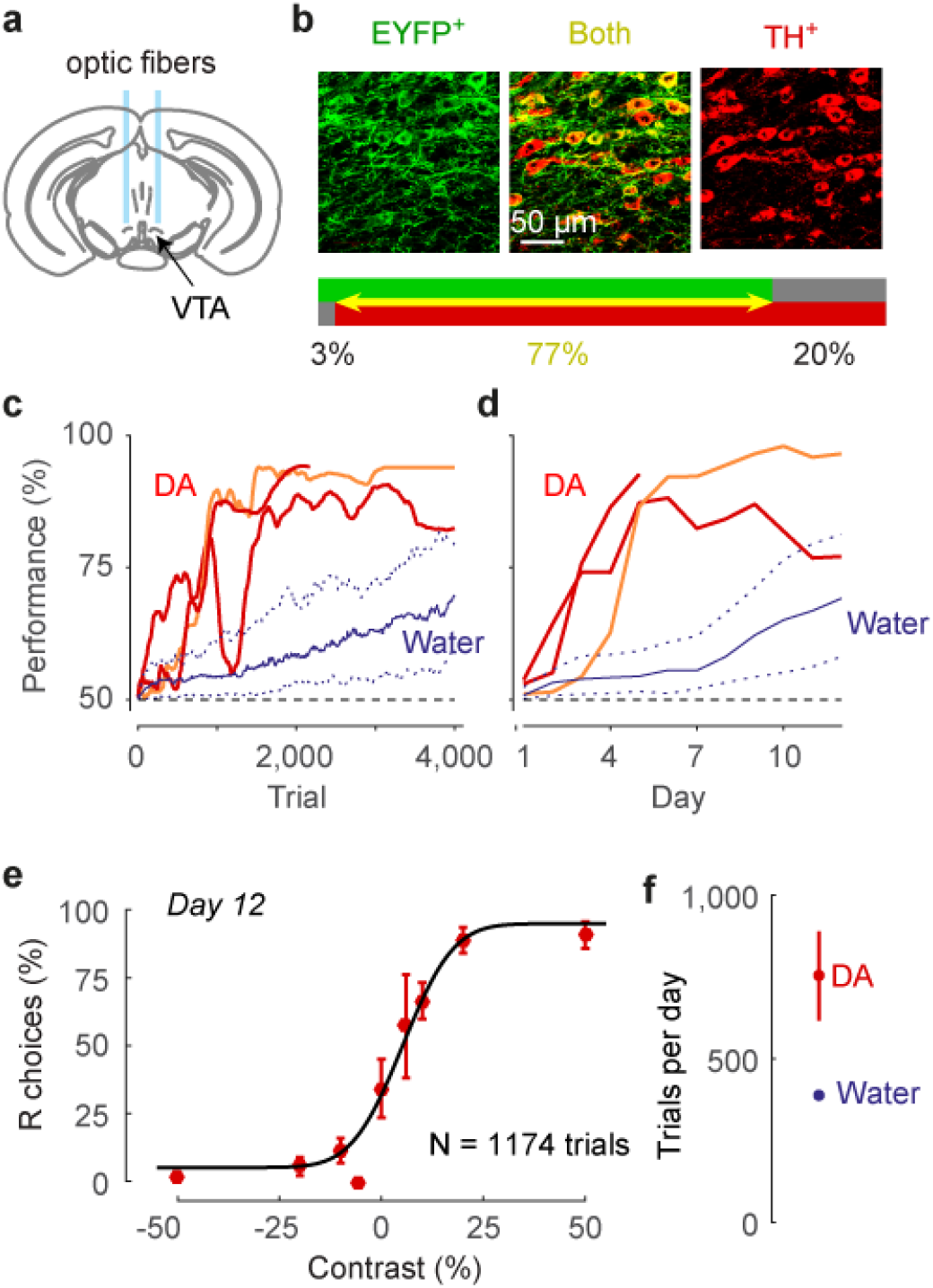
Using optogenetic phasic dopamine stimulation to train mice in the task. **a**. Schematic view of coronal section of the mouse brain (at Bregma -3.1 mm) showing ventral tegmental area (VTA), and fiber optics implanted above VTA to elicit release of dopamine (DA). **b**. Example images from a confocal microscope, showing expression of ChR2-EYFP (green) in TH+ (dopamine) neurons (red), and overlay showing both (yellow). The bars quantify the specificity of expression, showing statistics of ChR2-EYFP and TH+ expression in midbrain neurons (n = 1460 neurons counted in 121 confocal images acquired from 11 mice). **c,d**. Rapid learning of the task in three mice receiving DA stimulation as a reward. Red and orange lines show rapid increase in the performance of naïve mice that were trained solely with optogenetic DA stimulation. For comparison, blue curves show results for mice that trained with water reward (median and quartile ranges, replotted from Figure 1). **e**. Psychometric function obtained from example animal (orange line in **c,d**) on the 12^th^ day of behavioral training. Error bars show 95% binomial confidence intervals. **f**. Mean trials per day of mice receiving DA stimulation (red) compared to water rewards (blue). Error bars show s.e.m. (smaller than the dot for water rewards).

We then trained three naïve mice in our 2AFC task and reinforced correct choices with only optogenetic dopamine stimulation and an associated sound. In this preparation, a correct wheel response was followed by a short train (~ 0.5 s) of laser pulses, whose onset was marked by a click sound. The mice did not receive a water reward, and had free access to food and water in their home cage.

Mice trained with optogenetic DA stimulation rapidly learned the task, greatly outperforming mice trained for a water reward, both in speed of learning and in number of trials per session (Figure 5**c**-**f**). After only a few days of training with DA stimulation, mice often performed over 900 trials per session (in more than 50% of sessions), with high accuracy (>75%, Figure 5**c**-**d**), resulting in high-quality psychometric curves (Figure 5**e**). On average, mice rewarded with DA stimulation performed almost twice as many trials per session as those that were rewarded with water (Figure 5**f**). To assess the stability of DA stimulation as a means of providing reward, in one mouse we continued these measurements for 10 weeks, during which the method remained successful.

The click sound at the onset of the optogenetic stimulation may be important for the success of these experiments. We say this for two reasons. First, because of a pilot experiment where we attempted to train a mouse with optogenetic DA stimulation but no click sound, and the animal did not learn the task. Second, because it has long been known that sensory stimuli can be powerful secondary reinforcers (Herrnstein, 1964), and click sounds are particularly widespread and effective in “clicker training” (Pryor, 1999). We thus recommend pairing the optogenetic DA stimulation with a click sound.

### Stimulus discrimination

A method for performing psychophysics should be flexible, so that its design can be altered or made more complex as needed. For instance, the basic tasks that we have described, whether 2AFC or 2AUC, involve detecting whether a stimulus appears on the left or right side. To study mechanisms that combine information across hemispheres, however, it is useful to have the subject discriminate between stimuli that appear on both sides, as in contrast discrimination tasks that are commonly used with human observers (Boynton et al., 1999; Legge and Foley, 1980; Nachmias and Sansbury, 1974).

**Figure 6.**
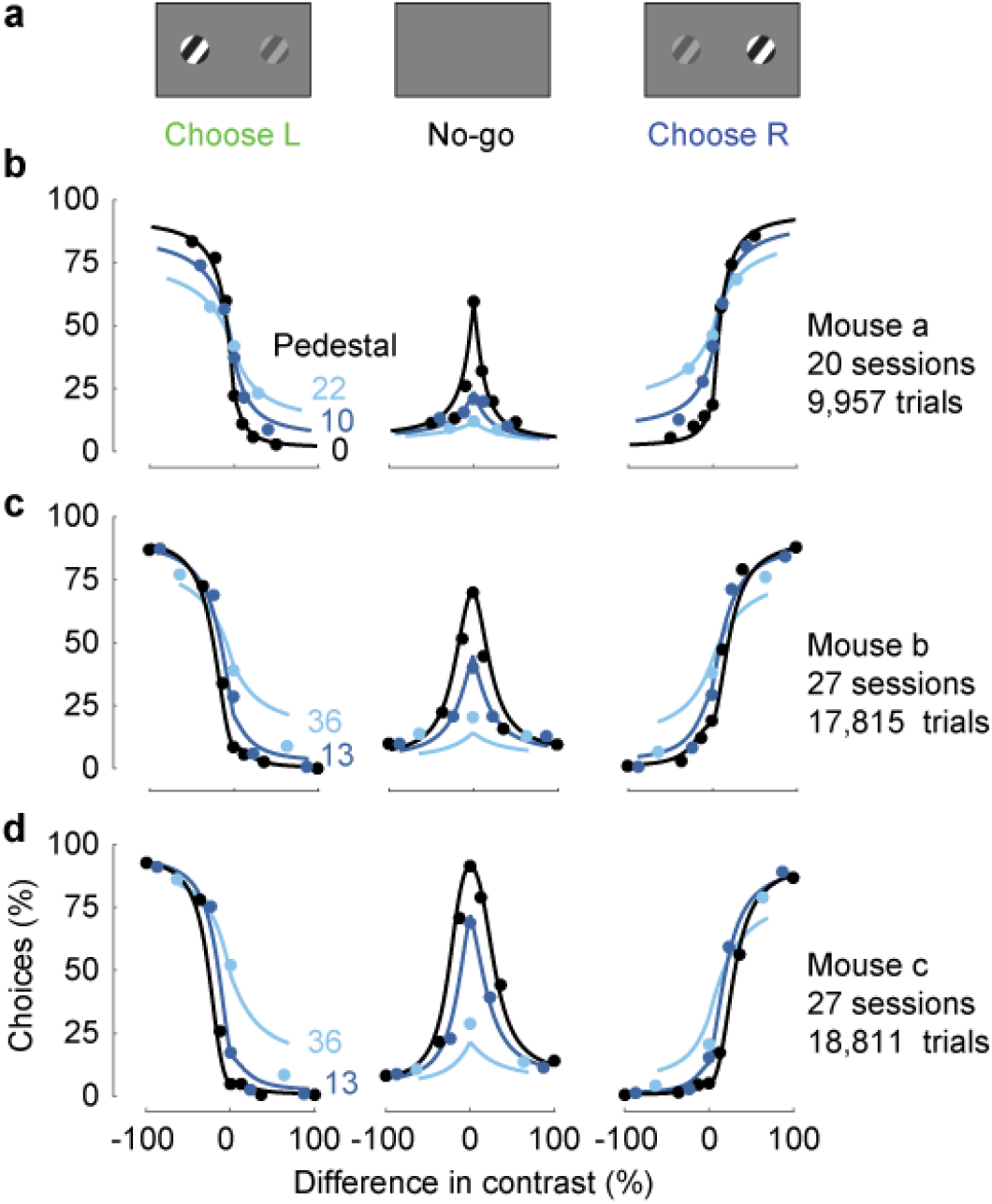
Extension of the 2AUC task to the study of contrast discrimination rather than detection. **a**. Schematic of the stimulus conditions used in the discrimination task. Gratings are presented on both sides and the mouse is rewarded for choosing the side with the highest contrast, or opting for no-go if both contrasts are zero. **b**. Psychometric data from 20 sessions in one mouse (9,957 trials). The three panels indicate, in the order, choices of the left stimulus, no-go, and choices of the right stimulus, as a function of the difference in contrast between the left and the right (c_R_-c_L_). The colors indicate the pedestal contrast, i.e. the minimum contrast present on the screen, min(c_L_,c_R_). **c,d**. Same as **b** for two more mice (27 sessions in each mouse, for a total of 17,815 trials in **c** and 18,811 trials in **d**).

Mice that had already learned the 2AUC detection task readily learned to perform a more general contrast discrimination task (Figure 6). In most trials, two stimuli appeared on the screen, and the mice were rewarded with water for selecting the stimulus that had the higher visual contrast (Figure 6**a**). A no-go response was rewarded only when no grating was presented on either side. If contrasts were nonzero and equal, mice were rewarded randomly with 50% probability for left or right responses. Mice learned this generalization of the task, yielding high-quality psychometric curves (Figure 6**b**). When one of the gratings had zero contrast (a “pedestal contrast” of 0%, Legge and Foley, 1980) the task was equivalent to contrast detection (e.g. Figure 3). When both gratings were present the mouse correctly gave fewer no-go responses, while finding it harder to indicate the side with higher contrast. Similar results were seen in all three mice (Figure 6**b**-**d**). Mouse decisions conformed closely to the predictions of the probabilistic observer model that we had introduced for contrast detection (Figure 3). With a fixed setting of its 6 parameters the model provided satisfactory fits to the 32 response probabilities measured across three pedestal contrasts.

These results illustrate the suitability of the task for studying sophisticated psychophysical paradigms. The task is flexible and can be modified in multiple ways, and it leads the mice to provide psychophysical curves that can be useful, in future studies, to measure the capabilities of mouse vision and to relate perceptual decisions to neural activity.

## Discussion

We have outlined a flexible task paradigm for assessing visual decision-making in mice. In this task, the mouse is head-fixed, allowing careful control of visual stimulation and measurement of eye position, and easing simultaneous brain recordings or manipulations. The steering wheel allows mice to accurately report one of two alternative stimuli, and the task is readily extended to allow a no response option (no-go response). The task is learned relatively quickly and highly reliably: most mice master it within a few weeks. The task yields a large number of trials within a single session, providing high-quality psychometric curves within individual sessions.

By designing a task that can be performed in a head-fixed context, we enhanced our ability to perform not only neural measurements, but also behavioral measurements such as tracking eye movements. While we found neuronal activity correlated with wheel turns, we saw that eye movements are sometimes executed during the same trial epochs as wheel turns, and may therefore also correlate with this activity. Tracking these behaviors and understanding the relationship of neural activity to both is therefore an important control as well as an interesting direction for further research.

The decisions that mice make in this task conform closely to the predictions of a simple probabilistic observer model. The model’s formulation in terms of multinomial logistic regression is closely related to an earlier formulation based on signal detection theory (Sridharan et al., 2014). Both formulations are two-dimensional, because their responses depend on the combination of two decision variables. As we have seen, this two-dimensional nature is essential to capture the effects of unilateral cortical inactivations. These effects would not be captured by models that account for multiple possible responses with a one-dimensional decision variable (Garcia-Perez and Alcala-Quintana, 2011, 2013; Kiani and Shadlen, 2009). Our formulation, moreover, has two advantages over the earlier one (Sridharan et al., 2014). The first advantage is technical: having a functional dependence on stimulus contrast minimizes the number of free parameters. The second has broader import: by recasting the model as a logistic regression it is easier to modify the analysis to include other predictor variables such as past history or neural activity.

Including past history or neural activity as additional predictors can be very useful. Indeed, it is common for animals and humans to be influenced by past decisions and outcomes even when such events are not informative (Abrahamyan et al., 2016; Bak et al., 2016; Busse et al., 2011; Licata et al., 2017). Moreover, including a neural signal as a predictor provides a means to assess whether that signal is informative of the animal’s choices: if adding the predictor improves the prediction of the choices, then the neural signal is informative as to the animal’s decisions (Nienborg and Macke, 2014).

We further demonstrated that transient optogenetic dopamine stimulation can substitute water reward. Transient dopamine stimulation had been shown to be sufficient to drive simple behavioral conditioning such as conditioned place preference (Tsai et al., 2009). Our results show that it is sufficient to motivate mice for making choices in a perceptual decision task. The combination of our task and dopamine stimulation may thus be useful for studying the effects of dopamine signals on perception and perceptual learning (Ding and Gold, 2013; Schultz et al., 1997).

Dopamine stimulation offers an attractive alternative to water control. It greatly accelerates task acquisition and almost doubles daily trial counts. Both advantages can be important. For instance, a large number of trials is particularly useful when relating perceptual decisions to neural activity. Moreover, the method is arguably less disruptive of the mouse’s normal behavior and physiology, as it does not involve controlling the animal’s water intake.

As currently implemented, however, our optogenetic method also carries limitations. First, the method requires the use of DAT-Cre mice, which may not be feasible if Cre needs to be expressed in other cells for other experimental purposes. Second, the method requires implantation of optic fibers, which take up valuable space on the head of the mouse. These limitations have prevented us, at this stage, from combining our dopamine stimulation preparation with 2-photon or widefield imaging methods.

An advantage of our behavioral task is that it is highly flexible, allowing for multiple extensions of the same basic design. We have modified the task depending on requirements, for example introducing a cue informing mice when to initiate their response, and a no-go response option to report stimulus absence. We exploited this no-go response in inactivation experiments, finding that inactivation of visual cortex diminished reports of contralateral stimuli but left ipsilateral reports unaffected. We also modified the task in a variant requiring contrast discrimination between two stimuli, generating high-quality psychometric functions that were modulated by contrast difference and by the level of the pedestal contrast. Further, we found that once trained, mice continue to perform if the stimulus position is fixed or is only transiently presented, which can be exploited to address concerns about stimulus movement being related to choice, or of presentation durations being controlled by the mouse.

We believe that the coupling of wheel movements to stimulus properties is a particularly useful learning aid, and is further generalizable. For instance, the task can be extended beyond the detection or discrimination of visual contrast. In preliminary results (not shown), we have trained mice to use the wheel to rotate a grating to a target orientation or to modulate the pitch of repetitively presented tones toward a target pitch.

Moreover, the continuous readout available from the steering wheel may provide further insight into the nature of behavior. We used the wheel to obtain binary or ternary reports, but the continuous readout may afford more sensitive behavioral assays, potentially probing factors such as motivation, confidence (Lak et al., 2014), response vigor, and vacillation (Resulaj et al., 2009). These considerations suggest extensions of the task to a fully interactive, flexible, and accurate platform to probe mouse vision and visuomotor behavior and establish their neural basis.

## Experimental Procedures

Experiments were conducted according to the UK Animals Scientific Procedures Act (1986) in male and female mice aged 8-24 weeks. To allow head-fixing, mice were first anesthetized and implanted with metal head-plate plate on the cranium. After at least 4 days recovery, mice were acclimatized daily for 3 days with handling and head-fixing. Mice were then trained in a simplified version of the task involving only stimuli with high contrast and no timing requirements. As performance improved, lower contrasts were gradually introduced and more stringent timings were required. Training criteria were qualitative and differed across experimenters and mice.

Most mice were trained using water as a reward: mice obtained water by performing the task. After performing the task, they received top-up fluids to achieve a minimum daily amount of 40 ml/kg/day. The weight in this calculation was measured before water control, and was updated daily based on a standard curve relating sex and age to body weight. Body weight and potential signs of dehydration were monitored daily.

Stimuli were presented on an LCD monitor or on 3 monitors placed around the animal. Intensity values were linearized with a photodiode. In some experiments, we attached plastic Fresnel lenses to the screens to make intensity spatially uniform. The response wheel was a Lego rubber tire, whose angle was measured using a rotary encoder. Water was dispensed by opening a solenoid valve. A detailed parts list is described at www.ucl.ac.uk/cortexlab/tools/wheel.

Stimuli were typically sinusoidal gratings in a Gaussian window. The specifics of this stimulus, however, generally differed across mice. To control visual stimulation, we used the Psychophysics Toolbox (Brainard, 1997; Pelli, 1997). To measure eye position and pupil dilation we used an infrared camera focused on one eye, illuminated by an infrared LED. For each video frame, we determined pupil size and location by fitting a 2D ellipse.

Imaging was performed in three 10-12 week old C57BL/6J female mice. During the initial surgery, we performed a craniotomy centered on the right primary visual cortex and injected a GCaMP6m virus under the human synapsin promoter (AAV2/1-*syn*-GCaMP6m-WPRE) (Chen et al., 2013). We then covered the craniotomy with coverslips and sealed it with dental cement. We began calcium imaging 3 weeks after virus injection. Imaging was performed using a Sutter two-photon movable objective microscope controlled by ScanImage (Pologruto et al., 2003), with excitation at 1,000 nm from a Coherent Chameleon laser, and focusing through an Olympus 20X objective. We chose a field of view with good GCaMP expression and mapped the preferred stimulus position of the field of view. This was chosen as the position of the task stimulus during behavior. We registered the raw calcium movies by aligning each frame to a reference frame (Guizar-Sicairos et al., 2008), and found cells through a semi-automated algorithm that selected nearby pixels that are significantly correlated with each other. We obtained a baseline *F_0_* by smoothing the calcium trace *F* in time and finding the minimum over a 20 s sliding window. We then computed Δ*F/F* by applying a causal exponentially weighted filter (τ = 0.2 s) to the fractional change *(F-F_0_)/F* (Jia et al., 2011).

To characterize psychometric performance in the 2-alternative forced-choice task (2AFC) we calculated the proportion of trials with rightward choices (ignoring repeat trials that were sometimes introduced after errors), and we fitted them with a standard psychometric function (e.g. Busse et al., 2011).

To measure task performance as a function of trial number we considered easy trials (contrast ≥ 40%) and estimated the probability of a correct response as a function of trial, and its confidence intervals (Smith et al., 2004). Daily performance was estimated by averaging across each day’s easy trials.

In the 2AUC version of the task the mouse was required to be still for 0.5-1 s after stimulus onset. This period of no movement was followed by an auditory Go cue. If the animal did not respond within 1.5 s of the Go cue, this was considered a No-go response. No-go responses were rewarded for trials with zero contrast stimuli or were met with a 2 s white noise burst for all other stimuli. We trained mice in this 2AUC version by first training them in the 2AFC version (at least with high contrast), and then introducing zero-contrast (No-go) trials.

To fit 2AUC data we used the probabilistic model in Equations 1-3. We fit the 4 parameters of Equation 2 through multinomial logistic regression, and optimized the two parameters in Equation 1. The resulting model has 6 parameters, but cross-validation indicated no loss in fit quality if one imposed *s*_*L*_ = *s*_*R*_, removing one parameter.

When measuring the effects of inactivation, we fitted the different inactivation conditions independently, while imposing that the parameters of Equation 1 were constant across conditions. This allowed us to capture the effects of inactivation with changes in the parameters of Equation 2.

Inactivation experiments were performed with transgenic mice expressing ChR2 in Pvalb-positive inhibitory interneurons, obtained by crossing a *Pvalb^tm1(cre)Arbr^* driver with an Ai32 reporter. Mice were prepared with a clear skull cap similar to that of Guo et al. (2014b) but with UV-curing optical adhesive. Light for inactivation was produced by a 473 nm diode laser coupled to a fiber, producing ~1.5 mW in a spot of ~0.3 mm diameter, positioned over visual cortex (3.3-3.7 mm posterior, 2.1 mm lateral) or somatosensory cortex (0.8 mm posterior, 2.5 mm lateral). Inactivation was performed randomly in ~30% of trials. Light was delivered as a 40 Hz sinusoid beginning 33.2±5.5ms (mean ± standard deviation) before visual stimulus onset and lasting until the mouse made a response. The task was 2AUC detection, but responses could be made immediately upon stimulus onset.

For optogenetic dopamine stimulation we used DAT-Cre mice (B6.SJLSlc6a3tm1.1(cre)Bkmn/J) backcrossed with C57/BL6J mice. We injected 1 µL of diluted virus (AAV5.EF1a.DIO.hChr2(H134R)-eYFP.WPRE) into VTA and SNc and implanted an optic fiber along the same track, with tip 0.5 mm above the injection site. We waited 3 weeks for virus expression, then started behavioral training. Upon making a correct choice, animals received a short train of laser stimulation (473 nm, 12 pulses each lasting 10 ms and separated by 40 ms, power 10-15 mW measured at the fiber tip) and a simultaneous click sound. Mice had free access to water.

To quantify efficiency and specificity of ChR2 expression in dopamine neurons, animals were anesthetized and perfused, brains were post-fixed, and 50 µm coronal sections were collected. Sections were then immunostained with antibodies to TH and EYFP and secondary antibodies labeled with Alexa Fluor 488 and 594 (Tsai et al., 2009). We quantified infection efficiency and specificity in 1,460 neurons from 121 confocal images collected from 11 mice.

The contrast discrimination task is based on the 2AUC task, but gratings could be presented on both sides of the screen simultaneously, and mice were rewarded for choosing the grating with the highest contrast (or rewarded 50% of the time if contrasts were equal). No-go responses were rewarded only if no stimulus was present. Mice were first trained in the 2AUC detection task, and discriminations were introduced incrementally in order of difficulty. Mice learned this discrimination task within a few days once they had learned the 2AUC detection task.

## Author Contributions

Conceptualization: CPB, AL, NAS, PZ-H, JFL, KDH, MC. Methodology: CPB, AL, NAS, PZ-H, SyS, SoS, JJP, MJW, LEW, KDH, MC. Software: CPB, AL, NAS, PZ-H. Formal Analysis: CPB, AL, NAS, PZ-H, MJW. Investigation: CPB, AL, NAS, PZ-H, CBR, EAKJ, AR, SyS, MJW. Writing – Original Draft: CPB, AL, NAS, PZ-H, MC. Writing – Reviewing and Editing: CPB, AL, NAS, PZ-H, EAKJ, JJP, KDH, MC. Funding Acquisition: CPB, AL, NAS, SyS, EAKJ, KDH, MC. Supervision: KDH and MC. Project Administration: MC.

## Acknowledgments

This work was supported by Senior Investigator Awards from the Wellcome Trust (MC and KDH), and by a Medical Research Council Doctoral Training Award (CPB), a Human Frontiers Science Program Fellowship (NS), a Sir Henry Wellcome Fellowship (AL), a Marie Skłodowska-Curie Fellowship (SS), and a Wellcome Trust PhD Studentship (EJ). JJP was funded by the Simons Fondation (SCGP 325476) and by the Champalimaud Foundation, and So.S. by the Fundação para Ciência e Tecnologia (SFRH/BD/51895/2012). MC holds the GlaxoSmithKline / Fight for Sight Chair in Visual Neuroscience.

---

**Supplementary Figure 1.**
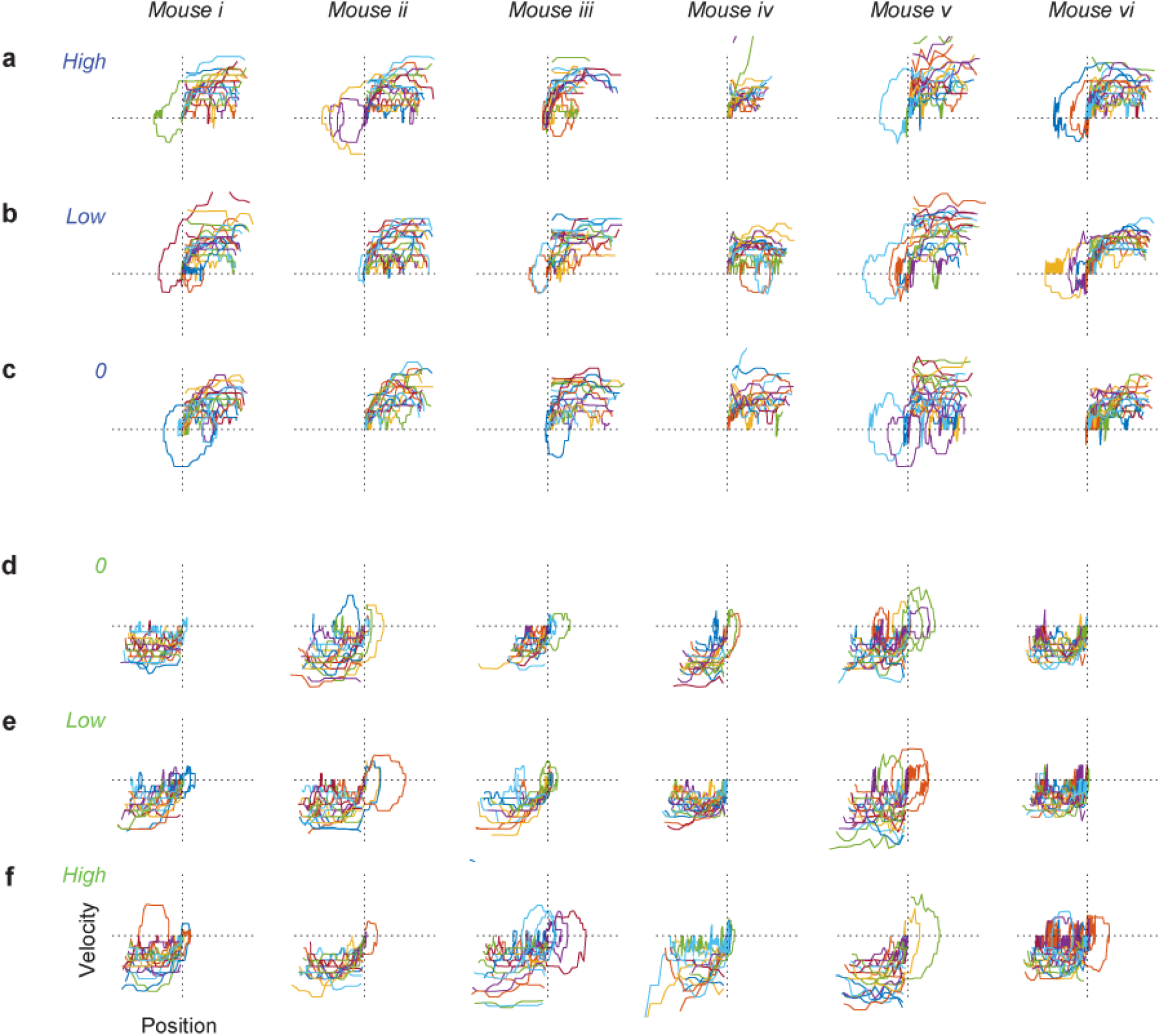
Trajectories of wheel turns made by mice in response to stimuli. Related to Figure 1. Traces show evolution of position and velocity during trajectories for turns made between stimulus onset and attainment of choice threshold. a-c: Trajectories that ended with a choice to the left, for stimuli that had high contrast on the left (a), low contrast on the left (b), or zero contrast (c). Any trials where the initial choice direction is inconsistent with the final choice must cross from one quadrant to the other (lower-left to upper-right), which is uncommon. d-f: Same as a-c, for trajectories that ended with a choice to the right.

**Supplementary Figure 2.**
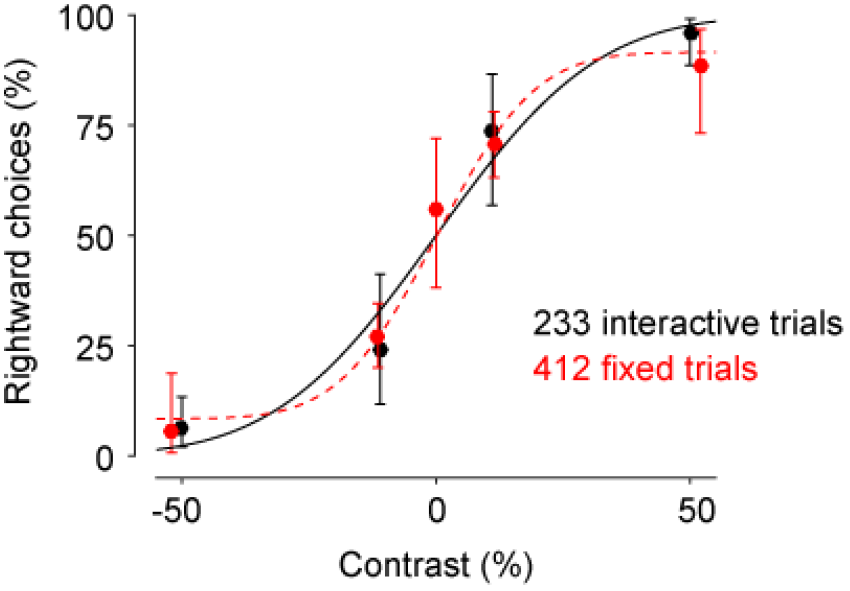
Comparison of psychophysical performance in interactive trials vs. fixed-stimulus trials. Related to Figure 1.These data were obtained in a single session in which two types of trial were randomly interleaved. In normal interactive trials, the steering wheel moved the stimulus (black). In the remaining trials, the mouse completes choices by turning the wheel as normal, but the stimulus remains fixed at the onset position (red). The ordinate plots the percentage of times the mouse chose the stimulus on the right (R), as a function of stimulus contrast (positive for R stimuli, negative for L stimuli). The psychometric curves fit across the two sets of trials (curves) are similar.

**Supplementary Figure 3.**
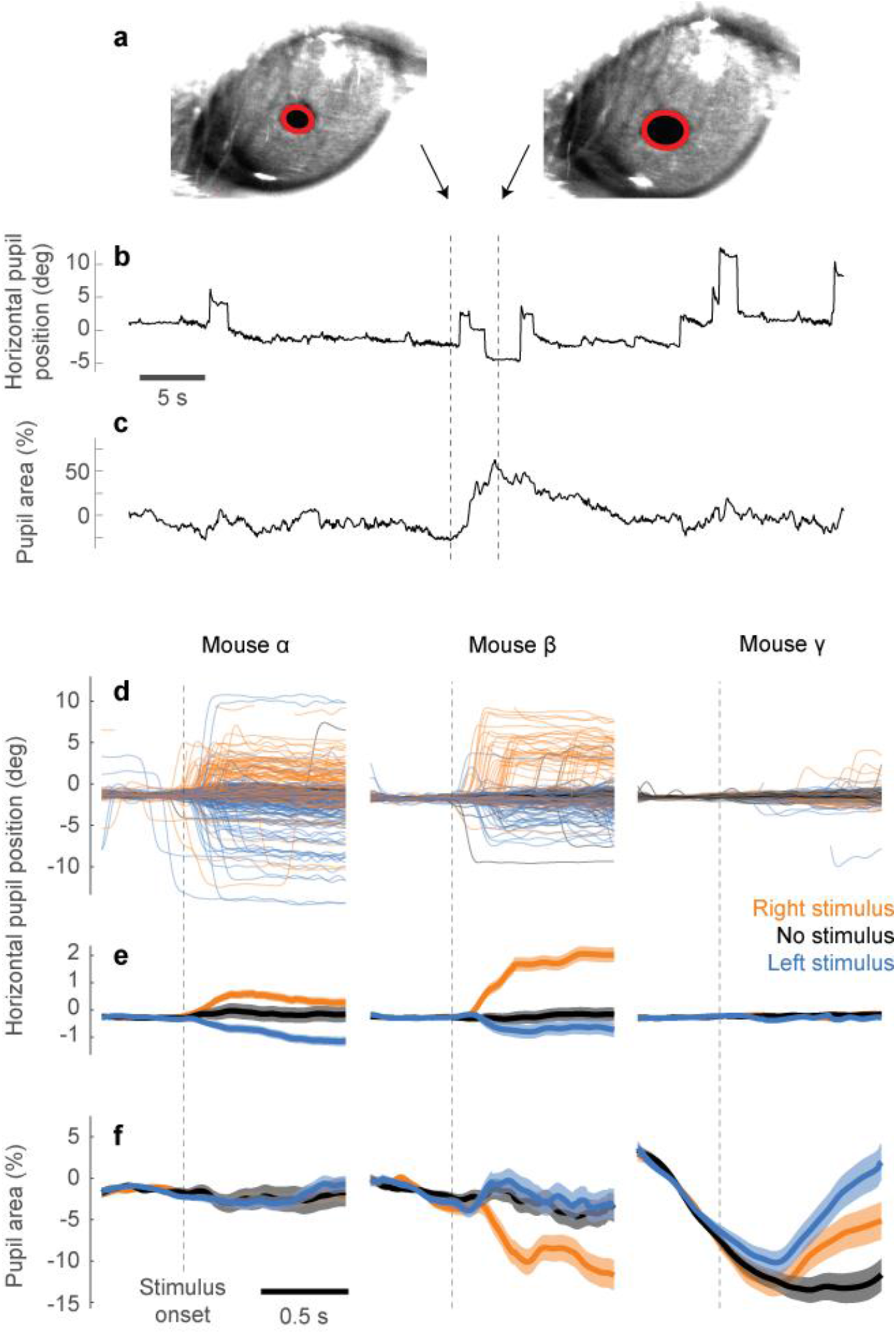
Eye movements during task performance. Related to Figure 1. a: Two example frames showing ellipses fit to the pupil. b: Example traces of horizontal position calculated from movies of the eye. Saccades as small as ~2 deg are clearly visible. Dashed lines indicate the times of the two frames in a. c: Same as b, for the pupil area (proportion change relative to the mean). d: Traces of pupil position for each trial from three example mice. Traces are aligned to stimulus onset and colored according to stimulus condition: stimulus on left (blue), right (orange), or no stimulus (black). e: Average of the traces in d. Notice different y-scale. Shaded area represents two s.e.m. f: Same as e, for the pupil area.

**Supplementary Figure 4.**
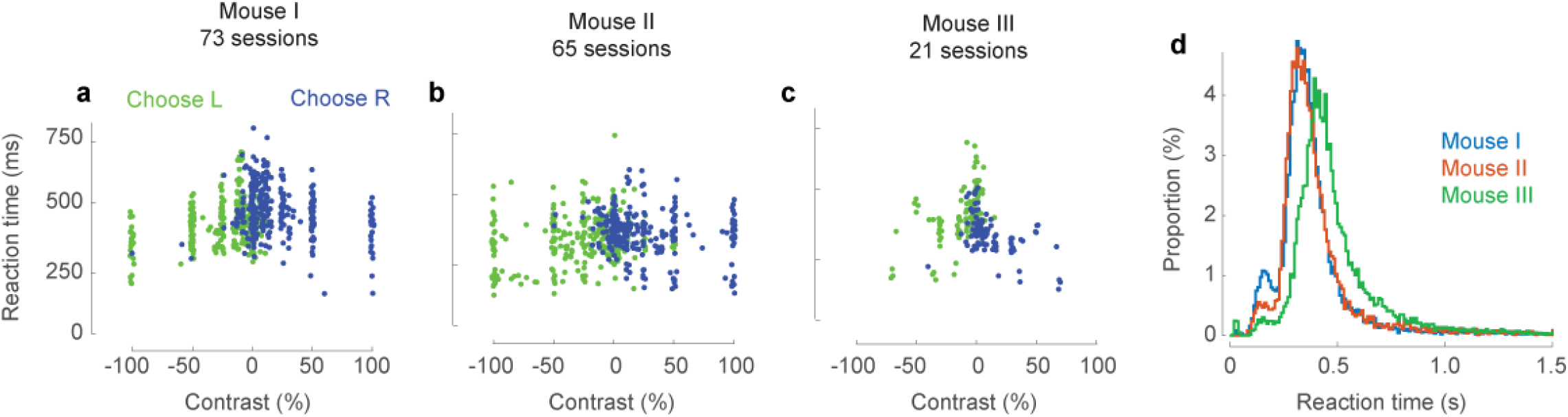
Reaction times in the 2AUC task. Related to Figure 3. a-c. Median reaction time for the example mice in Figure 3. Each dot indicates the median reaction time measured for stimuli of a given contrast in a single session, grouped by decisions made on the left (green) or on the right (blue). d: Distribution of reaction times for all trials in the three mice. The standard deviations of the reaction time distribution were 206, 181, and 198 ms respectively. The proportions of trials with reaction times longer than 1 s were 3.6%, 2.1%, and 2.8% respectively. The vast majority of reaction times, therefore, are much shorter than the 1,500 ms that would result in a No-go response.

**Supplementary Figure 5.**
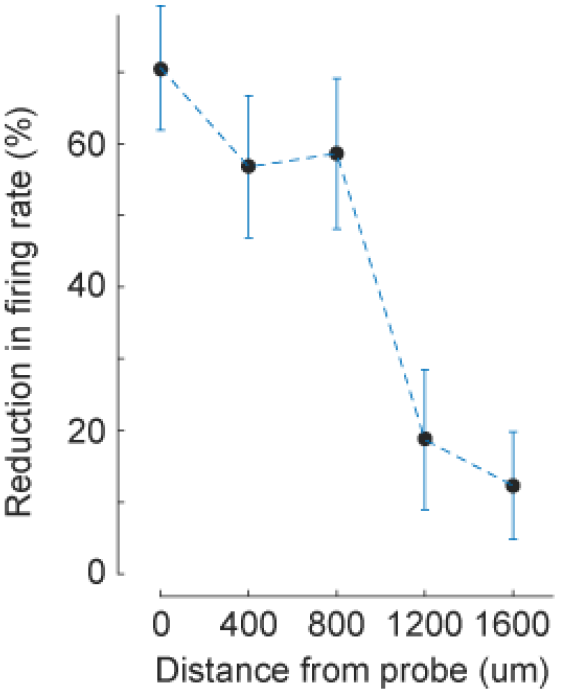
Control electrophysiological measurements show optogenetic inactivation of visual cortex was spatially focused, with a radius of ~1 mm. Related to Figure 4. We inserted custom multisite electrodes in visual cortex, and pooled responses from n = 110 single-unit and multiunit clusters with broad waveforms. We moved the laser at different distances from the electrode (abscissa) and measured the reduction in firing rate relative to control firing rate (modulation index, ordinate). The spot size and laser power (1.5 mW) were the same as in the behavioral experiments.

**Supplementary Figure 6.**
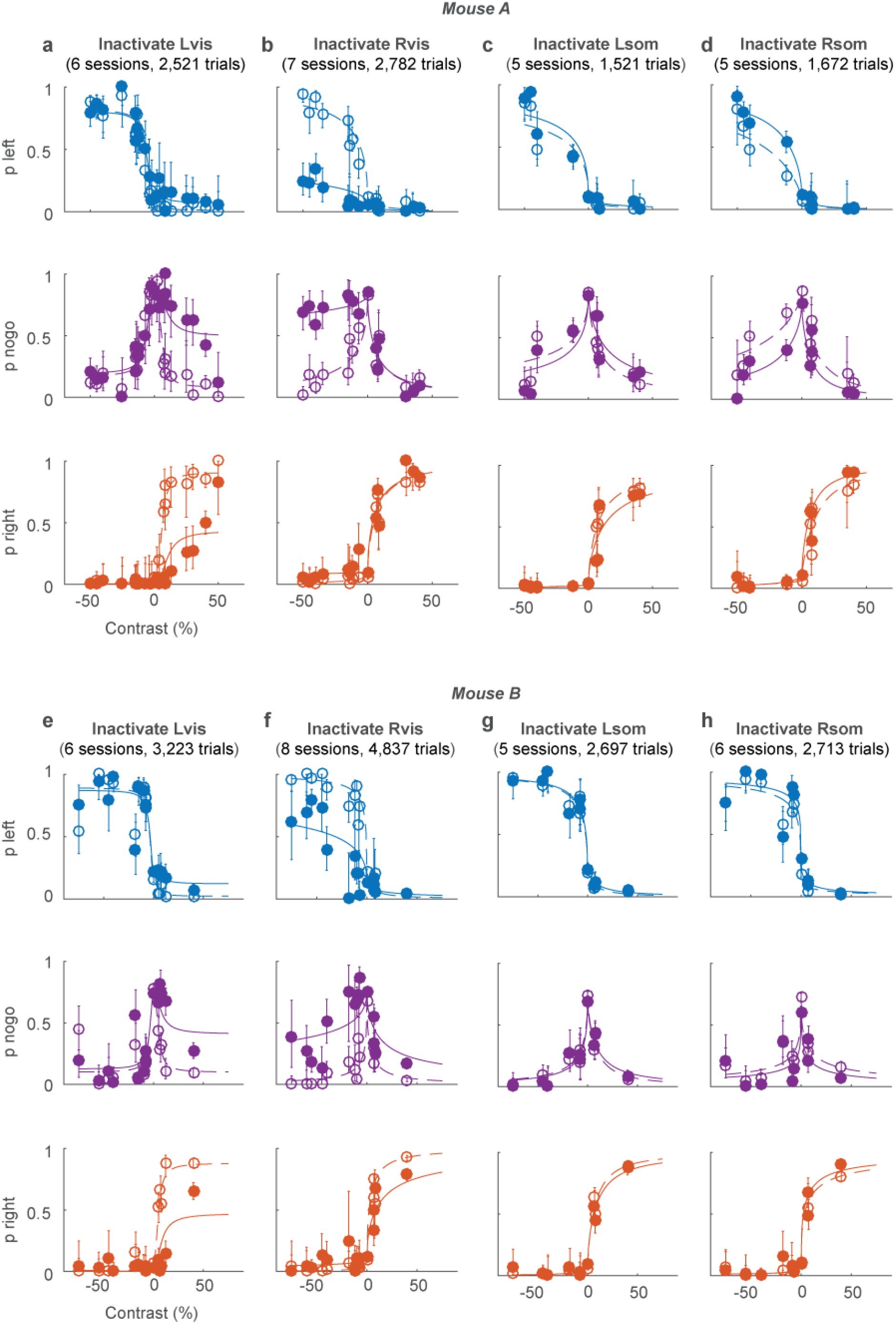
Results of inactivation in 4 regions in two mice. Related to Figure 4. **a,b**: Effects of inactivation of left and right visual cortex in mouse A (same data as Figure 4**b,c**). **c,d**: Effects of inactivation of left and right somatosensory cortex in mouse A. **e**-**h**: Same as **a**-**d**, for mouse B.

**Supplementary Figure 7.**
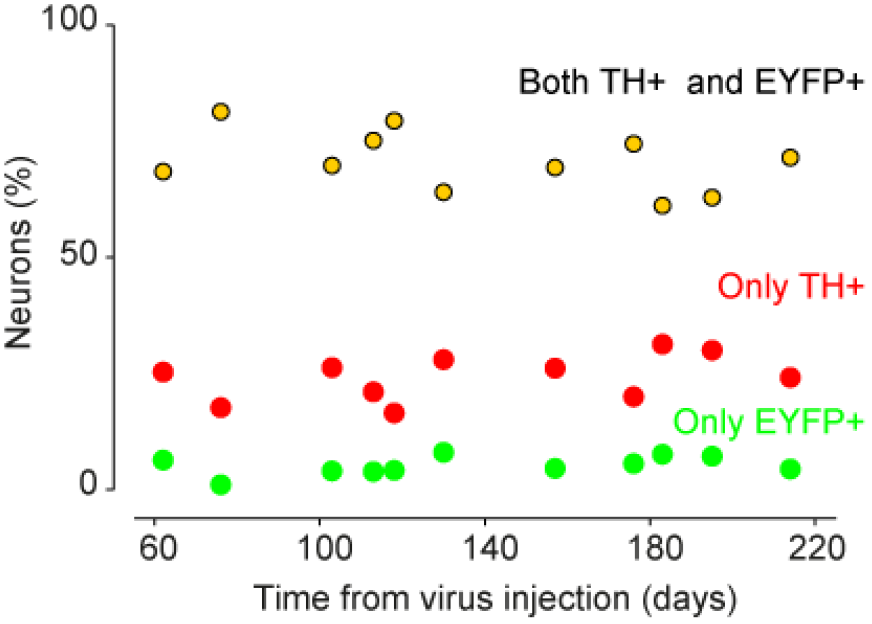
Stability of ChR2 expression in dopaminergic neurons. Data show measurements from 11 mice. Related to Figure 5.

